# Predicting *Mycobacterium abscessus* proteins with atypical amino acid composition essential for human infections

**DOI:** 10.1101/2025.03.24.645103

**Authors:** Guoqing Cheng, Ruth Howe, Binayak Rimal, Ngan Nguyen, Gyanu Lamichhane

**Author notes:** Equal contribution (Ruth Howe, Binayak Rimal). To whom correspondence should be addressed: Johns Hopkins University School of Medicine, 600 N Wolfe St, CMSC Room 3145, Baltimore, MD 21231.

## Abstract

*Mycobacterium abscessus* is an emerging opportunistic pathogen that causes chronic, difficult-to-treat lung infections, particularly in individuals with underlying lung disease or immune suppression. Despite its clinical significance, the fundamental biological systems of *M. abscessus* remain poorly understood due to limited research on this organism. Proteins that are unique to an organism are likely to contribute to the organism’s distinctive phenotypic traits. Therefore, we initiated this study by identifying proteins with unique features, hypothesizing that such proteins could be critical for *M. abscessus* pathogenesis. To identify these proteins, we analyzed the genome sequence of the laboratory reference strain using bioinformatics tools to detect proteins with unusual amino acid compositions. We then examined the genomes of a large collection of patient-derived *M. abscessus* isolates to predict proteins essential for the pathogen’s ability to cause disease in humans. Our analysis identified 10 proteins—MAB_0010, MAB_0039, MAB_1134, MAB_1602, MAB_1657, MAB_3052, MAB_3131, MAB_3413, MAB_4263, and MAB_4537—that exhibit restricted evolutionary variation in human infections, similar to five known essential proteins DnaA, DnaN, RpoA, RpoB and RpoC which comprise proteins involved in DNA and RNA synthesis. A majority of these proteins lack sequence homology with proteins of known function and are currently annotated as proteins of unknown function. The unique amino acid compositions of these proteins, their limited capacity to tolerate mutations, and their apparent exclusivity to *M. abscessus* suggest that they play essential roles in the pathogen’s ability to establish and maintain infection in humans. These findings highlight potentially promising targets for future drug development aimed at combating *M. abscessus* infections.

## INTRODUCTION

*Mycobacterium abscessus* (also known as *Mycobacteroides abscessus*) is an emerging opportunistic pathogen of growing concern in human health (1). Over the past few decades, the incidence of *M. abscessus* infections has been steadily rising worldwide (2, 3). Humans are believed to acquire this pathogen primarily from their immediate surroundings, as it is ubiquitous in aqueous environments, including public water systems and household plumbing (4–6). Its resilience and ability to thrive in diverse environmental niches make it particularly challenging to limit exposure to humans (7).

Phenotypically, *M. abscessus* exhibits several unique characteristics that distinguish it from other mycobacteria. It exists in either a rough or smooth colony morphology, with the ability to transition between these states without any detectable heritable genetic changes (8, 9). This morphological plasticity has been linked to its virulence and persistence in host tissues (10, 11). Additionally, *M. abscessus* is among the most antibiotic-resistant mycobacterial species, requiring combination therapy with no single-agent approach FDA-approved for treating this infection (12). Compounding the challenge of treatment, this pathogen readily forms robust biofilm, which enhances its resistance to antimicrobial agents, shields from host immune cells and contributes to its ability to cause chronic infections (13, 14).

Given that phenotypic traits are direct manifestations of genetic information, as outlined by the central dogma of molecular biology, we employed bioinformatics approaches to uncover unique genetic signatures in *M. abscessus* that may explain some of its distinct biological characteristics. The advent of high-throughput nucleic acid sequencing technologies has facilitated the rapid accumulation of whole-genome sequences for numerous organisms. In parallel, the development of powerful computational tools has enabled large-scale comparative genomic analyses, providing unprecedented opportunities to identify genetic determinants underlying specific phenotypes.

For this study, we sourced genome sequences of *M. abscessus* ATCC 19977 and a large collection of patient isolates from the NIH-NCBI GeneBank (15). This is a publicly available database curates pathogen genome sequences generated and submitted by independent laboratories. Because the archive is not restricted to a specific patient population or duration, it potentially represents a broad spectrum of *M. abscessus* infections across diverse demographics over several years.

Unlike *Mycobacterium tuberculosis*, which is transmitted exclusively between humans and has a restricted evolutionary niche, *M. abscessus* is primarily acquired from environmental sources (1, 2). This broader reservoir potentially provides increased opportunities for genetic diversification and adaptation. Additionally, *M. abscessus* has a significantly shorter doubling time (∼2 hours per cell cycle) compared to *M. tuberculosis* (∼20 hours per cell cycle), allowing it to evolve at a much faster rate (1). This rapid evolution, recorded in its genome sequences, makes large-scale genomic databases especially valuable for studying its adaptive mechanisms in the host from which they are derived.

Using bioinformatics methodologies, we systematically analyzed *M. abscessus* genome to identify proteins with unique amino acid compositions, sequences, and domains. We then assessed the potential requirement of these proteins in disease pathogenesis using a large collection of genomes. Since the genome sequences we analyzed were derived from patient isolates, they inherently represent genetic information that has successfully enabled *M. abscessus* to colonize human hosts (15).

Our approach was based on the hypothesis that proteins essential for survival in humans and, by extension, necessary to cause disease would exhibit evolutionary constraints, meaning they would sustain fewer mutations, particularly within functional domains essential for virulence as the genomes analyzed here were from patient-derived isolates and are presumed to represent a diseased state. To test this, we examined the extent and nature of mutations across different proteins, prioritizing those with minimal variation or predominantly synonymous mutations. To validate our findings, we included essential housekeeping proteins as controls, which are known to be highly conserved due to their fundamental cellular functions.

Through this analysis, we identified several proteins unique to *M. abscessus* and with unique amino acid sequence features that may play critical roles in its pathogenicity. We predict the proteins identified in this study that only sustain minimal mutations to be essential for *M. abscessus* to survive and cause disease in humans. These findings provide a valuable starting point for further functional investigations into the molecular mechanisms that contribute to the distinct and formidable nature of this pathogen. Understanding these unique genetic factors could ultimately inform the development of targeted therapeutic strategies to treat *M. abscessus* infections more effectively.

## RESULTS

### Proteins with atypical amino acid composition

To identify proteins with atypical amino acid compositions, it is essential to first establish the typical amino acid composition of proteins in *Mycobacterium abscessus*. This requires selecting a strain that is widely accepted as a reference for *M. abscessus*. For this study, we used strain ATCC 19977, the first recorded clinical isolate of *M. abscessus*, obtained in 1950 and archived at the American Type Culture Collection (ATCC) (16). Since ATCC 19977 was the first and the only *M. abscessus* isolate available from ATCC for decades, it has become the default laboratory reference strain for *M. abscessus* research. The genome of this strain has been sequenced, its genes and proteins have been annotated, and it has been included as a reference in genomic databases such as Mycobrowser and NCBI GenBank (15, 17). For this study, we obtained the genome and proteome sequences for ATCC 19977 from NCBI GenBank (RefSeq ID GCF_000069185.1) and used them as the basis for our analysis. We developed computational algorithms to perform the queries described below and used the ATCC 19977 genome and proteome data.

First, we analyzed the sequences of all proteins in ATCC 19977 to calculate the percentage distribution of each amino acid (**Figure S1**). This percentage distribution is referred to hereafter as the Global Proteome Amino Acid Distribution (**GPAAD**). The GPAAD of *M. abscessus* represents the expected mean percentage of each amino acid in a typical *M. abscessus* protein. Next, we calculated the percentage composition of each amino acid in individual proteins and compared the probability of this composition against the GPAAD for each protein (**Table S1**). Through this analysis, we identified *M. abscessus* proteins that contained specific amino acids at levels significantly deviating from the GPAAD for each amino acid (**Table 1**). Proteins with fewer than 100 amino acids were excluded from the analysis because the p-values for such proteins were often too low, leading to an overrepresentation of amino acid enrichment in these smaller proteins such as the ribosomal proteins.

**Table 1.** Proteins with atypical amino acid distribution are listed in this table. **Column 1**: Unique gene locus/protein ID. **Column 2**: Protein length in numbers of amino acids (AA). **Column 3**: Specific amino acid that the protein is enriched or depleted in. **Column 4**: % composition of the corresponding amino acid in the entire *M. abscessus* ATCC 19977 proteome (Global Proteome Amino Acid Distribution, GPAAD). **Column 5**: % composition of the corresponding amino acid in the corresponding protein in column 1. **Column 6**: Probability of observing the amino acid distribution relative to GPAAD.

| <u>Protein ID</u> | <u>Length (AA)</u> | <u>Amino acid</u> | <u>GPAAD %</u> | <u>AA% in Protein</u> | <u>Probability</u> |
| --- | --- | --- | --- | --- | --- |
| MAB_1303c | 214 | Alanine | 12.57 | 32.71 | 1.15E-14 |
| MAB_0108c | 201 | Alanine | 12.57 | 28.86 | 4.57E-10 |
| MAB_2522c | 79 | Cysteine | 0.78 | 11.39 | 1.30E-08 |
| MAB_2240 | 79 | Cysteine | 0.78 | 10.13 | 2.08E-07 |
| MAB_4932c | 182 | Aspartate | 6.00 | 20.33 | 4.61E-11 |
| MAB_4881c | 425 | Aspartate | 6.00 | 0.94 | 8.56E-08 |
| MAB_3052 | 227 | Aspartate | 6.00 | 0.44 | 1.16E-05 |
| MAB_4284c | 687 | Glycine | 8.79 | 23.87 | 2.72E-32 |
| MAB_0321c | 233 | Glycine | 8.79 | 35.19 | 5.49E-29 |
| MAB_1657c | 50 | Histidine | 2.23 | 28.00 | 3.07E-12 |
| MAB_4691c | 8108 | Lysine | 2.47 | 0.80 | 2.01E-29 |
| MAB_1314 | 238 | Leucine | 9.83 | 0.42 | 5.30E-10 |
| MAB_4263 | 932 | Methionine | 2.07 | 7.30 | 1.33E-18 |
| MAB_1134c | 981 | Methionine | 2.07 | 6.52 | 2.44E-15 |
| MAB_0630 | 638 | Proline | 5.72 | 16.46 | 2.54E-22 |
| MAB_2500 | 173 | Proline | 5.72 | 24.28 | 8.74E-16 |
| MAB_4810c | 803 | Glutamine | 3.21 | 11.58 | 4.91E-26 |
| MAB_2190 | 271 | Glutamine | 3.21 | 14.76 | 1.17E-15 |
| MAB_0978 | 744 | Arginine | 6.98 | 0.81 | 1.66E-16 |
| MAB_1013 | 792 | Arginine | 6.98 | 1.14 | 3.05E-15 |
| MAB_4284c | 687 | Arginine | 6.98 | 1.16 | 2.97E-13 |
| MAB_1084c | 108 | Serine | 5.74 | 27.78 | 2.51E-13 |
| MAB_1803 | 465 | Threonine | 6.06 | 16.13 | 1.01E-14 |
| MAB_3746c | 546 | Valine | 8.48 | 2.75 | 2.19E-08 |
| MAB_3413c | 712 | Tyrosine | 2.22 | 6.18 | 1.67E-09 |

For example, the protein encoded by gene locus *MAB_1303c* is composed of 33% alanine, while alanine constitutes only 13% of the GPAAD, making this protein the most enriched in alanine among all *M. abscessus* proteins. This protein, whose function remains unknown, is 214 amino acids long, with 70 of those residues being alanine. In contrast, the protein encoded by *MAB_1989c*, which is 189 amino acids long, contains only five alanine residues (3%).

For the proteins identified as having atypical amino acid compositions, we queried Mycobrowser (17) to determine their functional annotations based on homology to proteins with known functions or available functional descriptions. Of these proteins, 42.4% are categorized as proteins of unknown function or lacking homology to proteins with known functions. Consequently, the majority of *M. abscessus* proteins with atypical amino acid compositions remain functionally uncharacterized or lack detectable similarity to proteins with established functions (17).

### Proteins with domains containing atypical amino acid enrichment

In the analysis described above, the occurrence of an amino acid at a specific position within the primary structure of a protein was not considered relevant, as the objective was to identify proteins with atypical amino acid distribution across the entire protein. We hypothesized that considering the positional distribution of amino acids within a protein could reveal proteins in which specific amino acids are enriched in distinct linear regions. This could lead to the identification of primary structure domains enriched in particular amino acids. We propose that these regions, loosely referred to here as “domains,” may have evolved for specific cellular functions in *M. abscessus* that remain unknown.

To identify proteins with such features, we fragmented each protein’s sequence into 100-amino-acid segments and calculated the percentage composition of each amino acid within each fragment. Proteins with fewer than 100 amino acids were excluded from this analysis. This assessment identified 36 proteins enriched in at least one amino acid within a specific domain (**Table 2**). The proteins identified in this analysis were distinct from those identified in the previous analysis (**Table 1**).

**Table 2.**
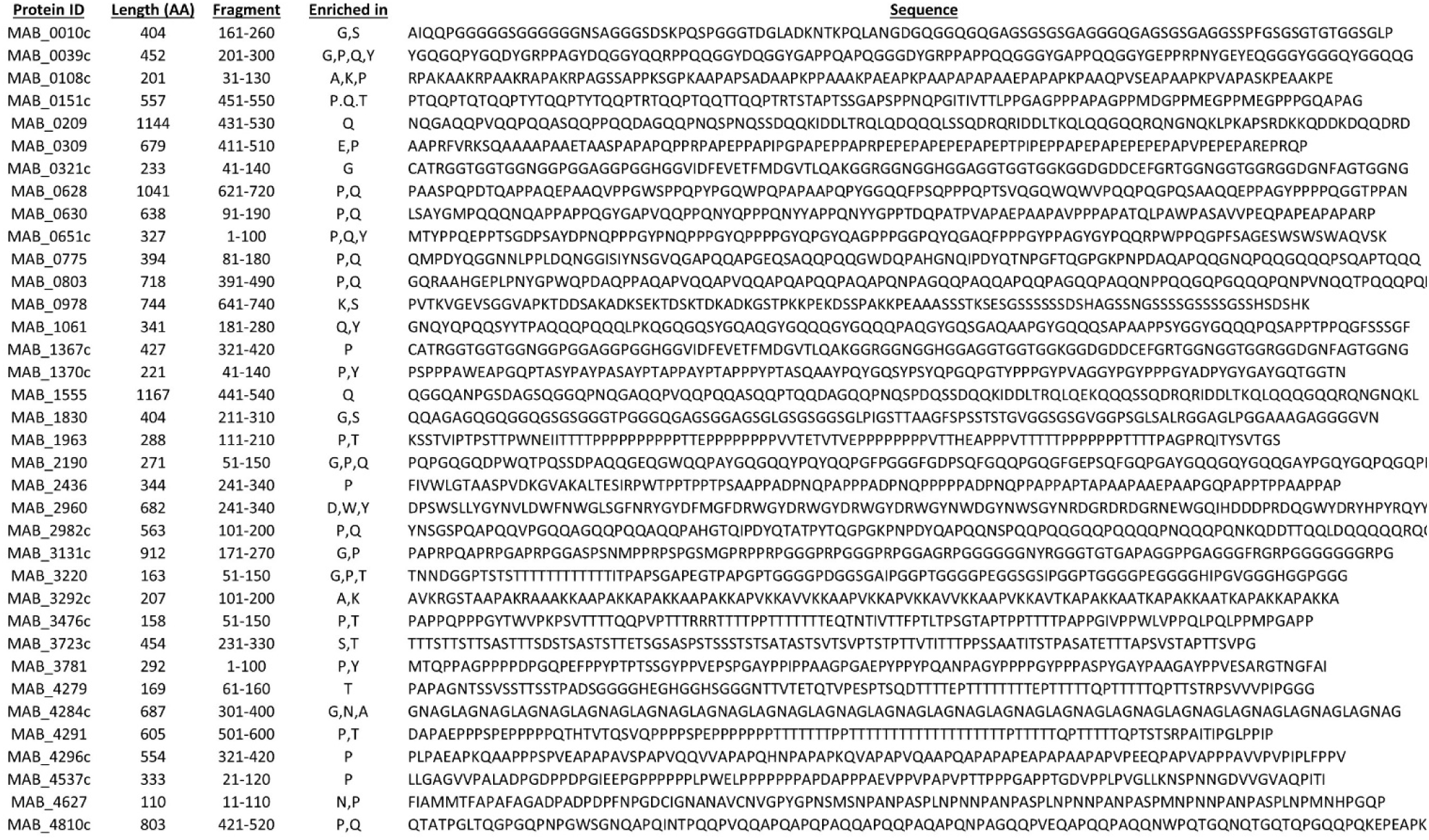
Proteins with domains containing atypical amino acid enrichment. **Column 1**: Unique gene locus/protein ID. **Column 2**: Protein length in number of amino acids (AA). **Column 3**: Fragment of the protein enriched in specific amino acid(s). **Column 4**: amino acid(s) that are enriched in the fragment. **Column 5**: Amino acid sequence of the corresponding fragment listed in the numerical order of the corresponding gene locus/protein ID.

#### Example 1: MAB_4291

MAB_4291 is a 605-amino-acid hypothetical protein with an unknown function. In the segment spanning positions 501–600, threonine constitutes 42% of the residues. This is notably higher than the 8.5% threonine content in the rest of the protein and the 6% typical threonine content in the general proteome of *M. abscessus,* GPAAD. This distinct enrichment suggests that the C-terminal threonine-rich region may serve a unique functional role in *M. abscessus*. Homologs of MAB_4291 were not identified in other mycobacteria, other bacteria, or higher organisms using BLASTP. DeepTMHMM (18) analysis predicts an extracellular N-terminus, one transmembrane domain spanning residues 457– 479, and an intracellular C-terminal threonine-enriched domain. Overall, MAB_4291 appears to be unique to *M. abscessus*, and its threonine-rich C-terminal region may contribute to a species-specific cellular function.

#### Example 2: MAB_0108c. (201 amino acids)

MAB_0108c is a 201-amino-acid protein with an unknown function. In the segment spanning residues 31–130, alanine and proline constitute 37% and 29%, respectively. This is markedly higher than the 13% alanine and 6% proline content in GPAAD. This enrichment in alanine and proline distinguishes MAB_0108c from a typical *M. abscessus* protein. BLASTP analysis did not identify any homologous proteins with known functions, and no homologs of this protein were found in other mycobacteria, other bacteria, or higher organisms.

#### Example 3: MAB_0978

MAB_0978 is a 744-amino-acid protein. In the segment spanning residues 641– 740, serine constitutes 33% of the amino acids, whereas it represents only 6% in GPAAD. While the overall serine content of MAB_0978 is 12%, the C-terminal end is distinctly enriched in serine. BLASTP analysis did not identify homologs of this protein in other mycobacteria or bacteria. Interestingly, the segment spanning residues 100–150 shares 45% amino acid identity with a NEPrilysin metallopeptidase in *C. elegans*. This suggests that MAB_0978 may have a specialized functional domain with evolutionary significance, potentially in interacting with the host.

This analysis revealed a subset of proteins in *M. abscessus* with unusual amino acid enrichment patterns in specific domains. These findings suggest that the positional distribution of amino acids may play a role in defining protein function and could provide insight into species-specific adaptations in *M. abscessus*.

### Proteins with Minimal Amino Acid Variety

The unique chemical properties of each amino acid enable proteins to carry out a wide range of biological functions. For example, cysteine residues can form disulfide bonds, which help stabilize a protein’s tertiary and quaternary structures. We assessed the amino acid variety in all *M. abscessus* proteins to identify those composed of the least variety of amino acids. Overall, 67.3% of *M. abscessus* proteins were composed of all 20 amino acids. Among the proteins with significantly restricted amino acid diversity, a few stand out (**Table 3**).

**Table 3.** Proteins with minimal amino acid variety. **Column 1**: Unique gene locus/protein ID listed in numerical order. **Column 2**: Protein length in numbers of amino acids (AA). **Column 3**: ‘variety’ refers to the number of different amino acids present in the protein. **Column 4**: Amino acids (one letter alphabet code) missing in the protein. **Column 5**: Probability of observing the amino acid distribution relative to GPAAD.

| **Protein ID** | **Length (AA)** | **Variety** | **Missing AAs** | **Probability** |
| --- | --- | --- | --- | --- |
| MAB_0030 | 65 | 10 | C,D,F,H,I,K,N,T,W,Y | 9.23E-10 |
| MAB_0108c | 201 | 15 | C,F,H,W,Y | 1.99E-09 |
| MAB_0169c | 271 | 16 | C,H,W,Y | 9.56E-09 |
| MAB_0319 | 114 | 14 | C,F,H,N,W,Y | 7.40E-07 |
| MAB_0448c | 67 | 14 | D,E,K,Q,S,W | 6.10E-08 |
| MAB_0672c | 249 | 16 | C,N,W,Y | 2.86E-08 |
| MAB_1602 | 204 | 15 | C,H,R,W,Y | 3.53E-13 |
| MAB_4074c | 145 | 17 | H,R,W | 1.14E-07 |
| MAB_4592c | 159 | 16 | C,D,H,N | 9.24E-09 |
| MAB_4774c | 115 | 16 | D,F,H,Q | 3.83E-08 |
| MAB_5000c | 37 | 13 | A,E,F,L,T,W,Y | 1.56E-07 |

#### Example 1: MAB_1602

MAB_1602 is a 204-amino-acid protein that lacks cysteine, histidine, arginine, tryptophan, and tyrosine — which represent approximately 1%, 2%, 7%, 2%, and 2%, respectively, of the total amino acid content in the *M. abscessus* GPAAD. The protein’s function remains unknown. DeepTMHMM predicted it to be a secreted protein. Homologs of MAB_1602 are found only in *Mycobacterium* species, suggesting that it may play a role specific to mycobacterial physiology.

#### Example 2: MAB_0448

MAB_0448 is a short 67-amino-acid protein that lacks six amino acids: aspartate, glutamate, lysine, glutamine, serine, and tryptophan. The likelihood of lacking these specific residues, given their general representation in the *M. abscessus* proteome, is calculated to be **6.1 × 10⁻⁸** — indicating that this is statistically improbable and likely the result of selective evolutionary pressure. Homologs of MAB_0448 could not be identified in other mycobacteria, other bacteria, or any other known organisms, suggesting that it may have evolved *de novo* relatively recently to meet a species-specific functional need. Its restricted amino acid composition could confer a selective advantage in *M. abscessus’s* environmental niche.

Despite its short length, DeepTMHMM predicts two transmembrane domains spanning residues 2–24 and 28–50 with over 80% probability. The presence of these transmembrane domains in a protein unique to *M. abscessus* suggests that it may play a structural role in stabilizing the cell wall or contributing to membrane function under environmental stress, such as its aqueous habitats where fluctuations in temperature, salinity, and pH are common.

#### Example 3: MAB_0030

MAB_0030 is a 65-amino-acid protein and has the least amino acid variety of any protein in the *M. abscessus* proteome. It lacks 10 amino acids: cysteine, aspartate, phenylalanine, histidine, isoleucine, lysine, asparagine, threonine, tryptophan, and tyrosine. The probability of lacking these specific residues, based on their overall representation in the *M. abscessus* proteome, is 9.2 × 10⁻¹⁰, making this an extremely rare amino acid composition. The function of MAB_0030 remains unknown. Interestingly, BLASTP revealed that homologs of this protein have been identified in several non-mycobacterial species, including the yet-to-be-cultured *Candidatus Brocadia sinica*. This suggests that while MAB_0030 may have originated in a non-mycobacterial ancestor, it has undergone significant divergence and specialization in *M. abscessus*.

### Evolution of *M. abscessus* Proteins with Atypical Amino Acid and Identification of Proteins Essential for Causing Disease in Humans

*M. abscessus* is known to persist in diverse aqueous environments, ranging from natural water bodies (e.g., rivers, lakes) to man-made water systems, including household plumbing and hospital water supplies (4, 5). These environments serve as reservoirs from which humans are frequently exposed to and potentially infected by *M. abscessus* through inhalation or direct contact. Indeed, several *M. abscessus* outbreaks have been linked to hospital water sources and contaminated equipment, exposing patients to the likely most human-adapted strains (19–21). This environmental persistence underscores the importance of *M. abscessus*’s ability to adapt to a wide range of ecological niches, which likely contributes to its extensive genotypic and phenotypic diversity.

### Proposed Model of *Mab* Evolution in Humans

Given the evidence of *M. abscessus*’s diverse environmental origins and its capacity to cause lung disease, we propose a model to describe the evolutionary pathway of pathogenic *M. abscessus* strains (**Figure 1**). According to this model, humans are exposed to a heterogeneous population of *M. abscessus* strains through inhalation/contact. Within the host environment, selective pressures drive the evolution of specific strains capable of surviving and establishing infection in the lung. Pathogenic strains are subsequently isolated in clinical settings, where whole-genome sequencing enables comparative genomic analysis. This analysis allows for the identification of genetic adaptations that may facilitate infection and persistence in the human host. Figure 1 illustrates how different *M. abscessus* genotypes are acquired by different individuals, resulting in clinical infections. By comparing the amino acid sequences of key proteins across these clinical isolates, we aim to identify evolutionary patterns that reflect selective pressures within the host environment and identify proteins essential for establishing disease in humans.

**Figure 1.**
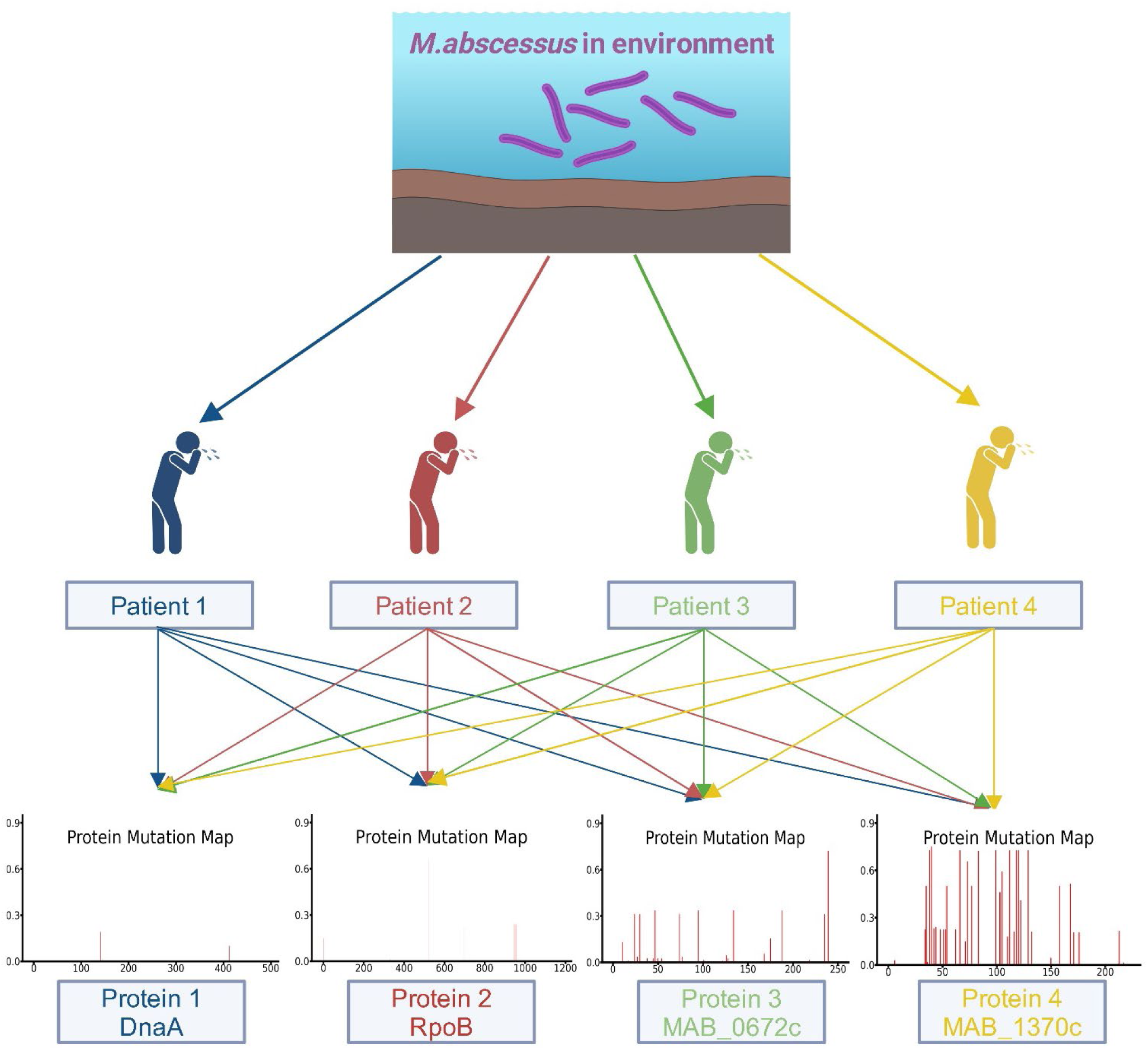
A model describing the natural history of *M. abscessus* infection in humans and the selection of genotypes that cause disease. This model illustrates the acquisition of genetically diverse *M. abscessus* isolates from human dwellings. It includes an analysis of amino acid sequences, highlighting specific mutations and their frequencies in patient-derived isolates. To illustrate the evolutionary pressures on essential proteins, amino acid mutation map for the DNA replication initiation protein (DnaA) and the RNA polymerase β-chain (RpoB) in clinical isolates are shown. These proteins exhibit low mutation frequencies, consistent with their essential role in *M. abscessus* survival. In contrast, two other proteins with distinct mutation profiles are also shown. These proteins harbor a high number of amino acid mutations, suggesting that they are under less selective pressure and therefore likely not essential for survival in humans.

### Identification of Essential Proteins Using Comparative Genomics

To implement this model, we analyzed publicly available genome sequences from 430 clinical *M. abscessus* isolates. We first determined mutation profile in five proteins known to be essential for DNA and RNA synthesis to use a comparator for essentiality of other proteins. The selected proteins include: chromosomal DNA replication initiation protein, DnaA (MAB_0001), DNA polymerase III β-subunit, DnaN (MAB_0002), RNA polymerase α-chain, RpoA (MAB_3770), RNA polymerase β-chain, RpoB (MAB_3869), RNA polymerase β’-chain, RpoC (MAB_3868).

We mapped and quantified both nucleotide and amino acid mutations in these proteins, calculating the ratio of amino acid mutations to nucleotide mutations (**A/N index**) (**Table S2**). A lower A/N index indicates that most nucleotide mutations are silent (synonymous), which is expected for proteins that are essential for basic cellular function. In essential proteins, amino acid substitutions are often detrimental, leading to reduced fitness or loss of viability.

Our analysis showed that the mean A/N index for the five essential proteins was 0.043, indicating minimal capacity for amino acid substitution to maintain functional integrity. This observation is consistent with prior studies indicating that essential proteins involved in fundamental cellular processes are highly conserved and resistant to amino acid changes (22). For example, in DnaA and RpoB, the majority of nucleotide mutations were synonymous, reinforcing the essentiality of these proteins (**Figure 2A** and **2B**).

**Figure 2.**
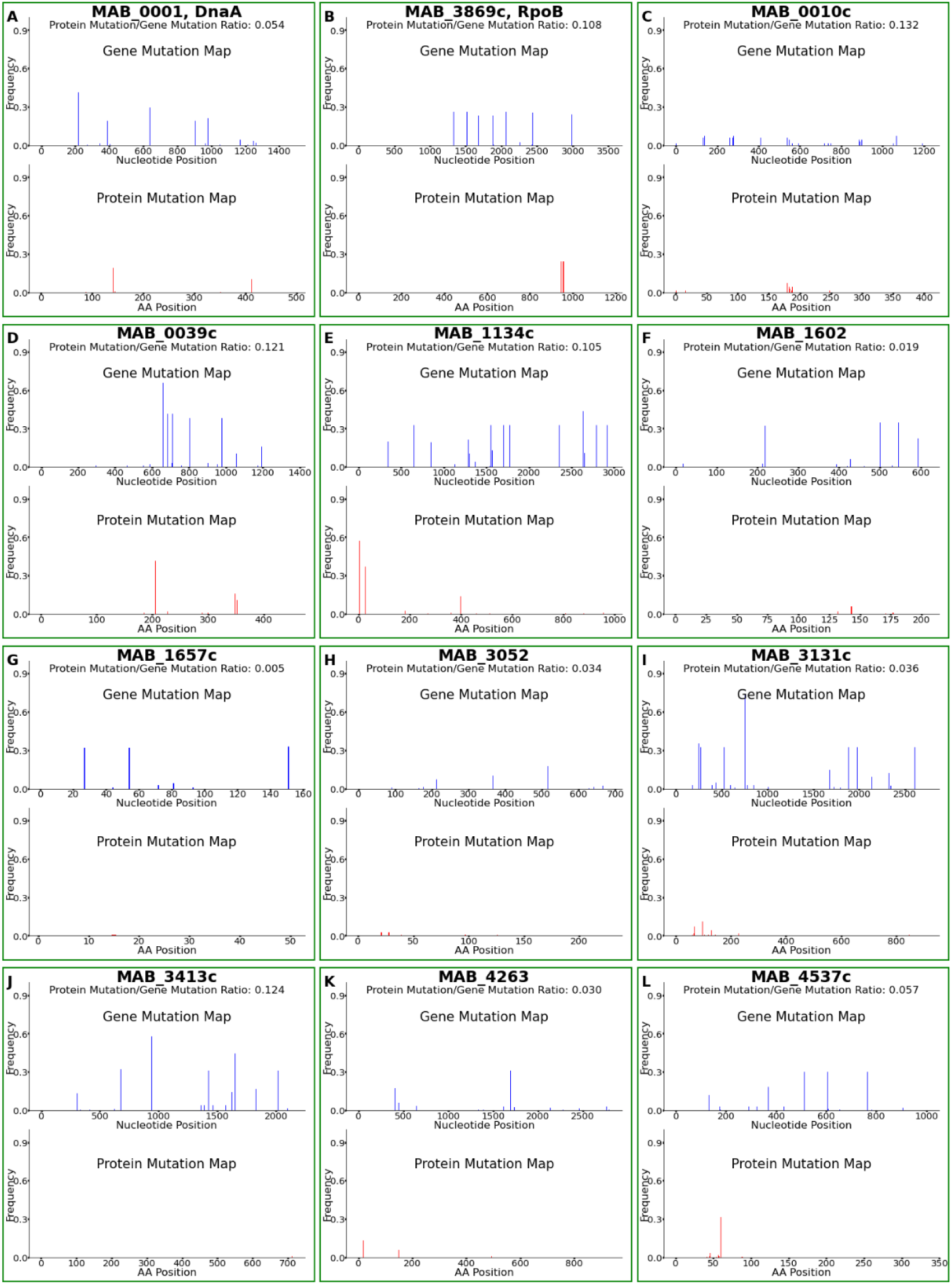
Mutation maps that illustrate the evolution of *Mab* genes/proteins. Each panel shows the mutation profile of a specific gene and corresponding protein, highlighting the nucleotide and amino acid positions and the frequency of mutations in patient-derived isolates. Only genes and corresponding proteins with amino acid mutation frequencies similar to known essential proteins, such as DnaA (panel A) and RpoB (panel B), are shown. Mutation maps for proteins with high amino acid mutation rates are provided in Table S3, Figure S2.

### Predicting Novel Essential Proteins

Using the *M. abscessus* A/N index, we extended this analysis to identify other potentially essential proteins in *M. abscessus*. Among the 63 proteins with atypical amino acid distributions identified above, 10 proteins exhibited an A/N index similar to that of the five known essential proteins (**Table 4**, **Figure 2**). These proteins, which this analysis predicts to be essential for *Mab* survival and pathogenesis, include: MAB_0010, MAB_0039, MAB_1134, MAB_1602, MAB_1657, MAB_3052, MAB_3131, MAB_3413, MAB_4263 and MAB_4537.

**Table 4.**
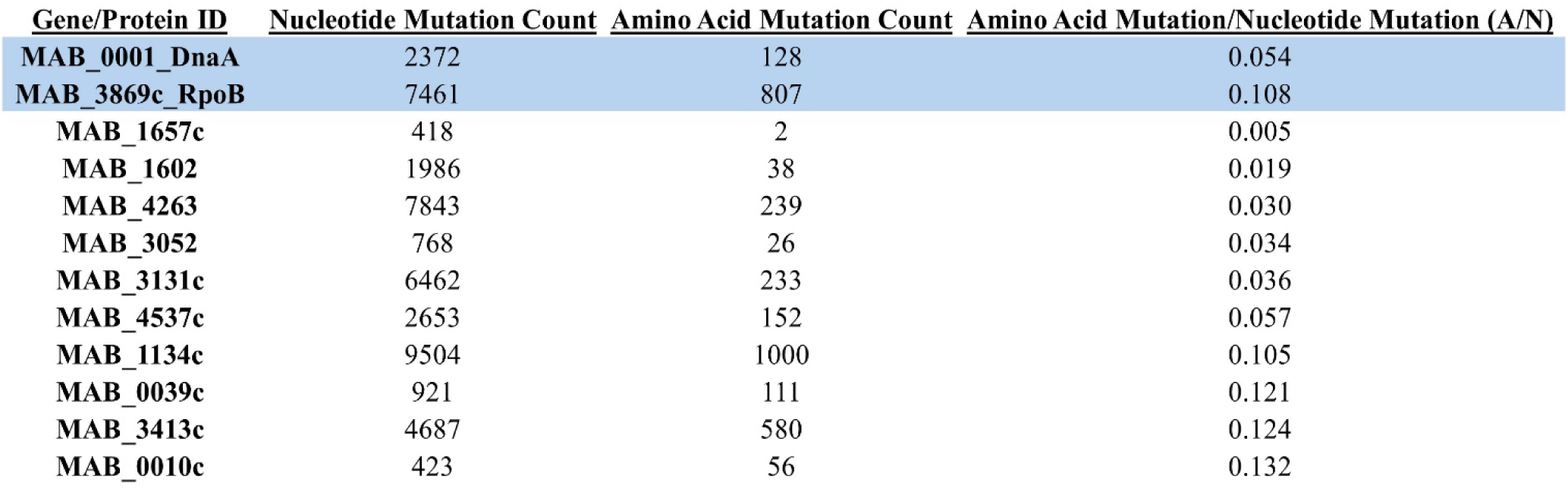
Predicted *M. abscessus* Proteins Essential for Disease Pathogenesis in Humans. The first two proteins—chromosomal DNA replication initiation protein (DnaA) and RNA polymerase β-chain (RpoB)—are well-established as essential for cell survival and are included here for comparison. The number of nucleotide and amino acid mutations identified in the corresponding genes and proteins from patient-derived isolates are shown in columns 2 and 3, respectively. Column 4 lists the amino acid mutation-to-nucleotide mutation (A/N) ratio for each gene.

| Gene/Protein ID | Nucleotide Mutation Count | Amino Acid Mutation Count | Amino Acid Mutation/Nucleotide Mutation (A/N) |
| --- | --- | --- | --- |
| MAB_0001_DnaA | 2372 | 128 | 0.054 |
| MAB_3869c_RpoB | 7461 | 807 | 0.108 |
| MAB_1657c | 418 | 2 | 0.005 |
| MAB_1602 | 1986 | 38 | 0.019 |
| MAB_4263 | 7843 | 239 | 0.030 |
| MAB_3052 | 768 | 26 | 0.034 |
| MAB_3131c | 6462 | 233 | 0.036 |
| MAB_4537c | 2653 | 152 | 0.057 |
| MAB_1134c | 9504 | 1000 | 0.105 |
| MAB_0039c | 921 | 111 | 0.121 |
| MAB_3413c | 4687 | 580 | 0.124 |
| MAB_0010c | 423 | 56 | 0.132 |

MAB_1134 and MAB_4263 which encode for transmembrane channels belonging to the Mycobacterial Membrane Protein Large (MmpL) that are involved in lipid transport (23). Several MmpL homologs in *M. tuberculosis* are essential for survival and to cause infection in animal models (24). MmpL3 has been pursued as a target for drug development against *M. tuberculosis* and *M. abscessus* (25–27). However, as a majority of the proteins identified in this assessment lack homologs in other bacteria, it suggests that their functions may be highly specific to *M. abscessus*’s adaptation and survival in the human host environment. The lack of known homologs implies that these proteins could represent unique targets for therapeutic intervention or biomarkers for infection.

## DISCUSSION

The incidence of diseases caused by non-tuberculous mycobacteria (NTM) has now surpassed that of tuberculosis in the United States (28). *M. abscessus* is the second most commonly isolated NTM from patients and is regarded as the most difficult to treat due to the lack of FDA-approved treatments for this pathogen (12, 29). Current treatment strategies rely on repurposed drugs and expert opinion rather than direct evidence from randomized clinical trials (30–32). Cure rates with existing therapies remain low, ranging between 30% and 50%, and resistance to these antibiotics is increasing (33). Consequently, there is an urgent need for more effective drugs to treat *M. abscessus* infections.

A promising strategy for drug discovery involves identifying proteins that are unique to *M. abscessus* and essential for its ability to cause disease. This approach has been successful in identifying potential drug targets in other mycobacteria while also providing insight into the genes and proteins essential for survival and pathogenesis (24, 34–37). Despite the increasing incidence of *M. abscessus* infections, research into its basic biology has lagged behind that of related mycobacteria, such as *M. tuberculosis*. However, recent advances in computational biology and the availability of large genomic datasets have made it possible to explore the fundamental biological systems of *M. abscessus* in greater detail.

This study aimed to identify proteins with atypical amino acid composition that are unique to *M. abscessus* and predict which proteins are potentially essential for its ability to cause disease in humans. The *M. abscessus* complex consists of three subspecies: *abscessus*, *massiliense* and *bolletii* (38). To identify these proteins, we used the annotated genome of the *M. abscessus* reference strain ATCC 19977, which belongs to the subspecies *abscessus*, the most commonly seen in clinical infections (2). The gene and protein annotations provided by Mycobrowser (17), a widely used reference for *M. abscessus*, were used to develop our computational algorithms.

Computational algorithms were developed and executed to query the genome data for proteins with atypical amino acid compositions. The algorithm outputs were manually verified, and adjustments were made to ensure accurate data assessment. Statistical analysis was then performed on the results, and only proteins that met strict statistical criteria were reported as top hits. The identified proteins are listed in Tables 1, 2, and 3, while additional proteins that met these criteria are provided in the Supplementary Information.

Identifying essential proteins is critical for understanding the molecular basis of *M. abscessus* infections and for developing targeted drug therapies. Recently, two independent studies identified *M. abscessus* genes essential for *in vitro* growth using a transposon mutagenesis approach (39, 40). While these studies identified genes essential for growth *in vitro*, they did not capture genes required for disease pathogenesis in humans. In previous studies on related mycobacterium, transposon mutants were used to infect animal models to identify genes essential for disease establishment (24, 36, 37). However, differences between human and animal immune systems mean that all of these findings may not translate directly to human infections.

To address this limitation, we used a large collection of *M. abscessus* isolates derived from human patients to predict genes and proteins essential for disease in humans. Since these isolates represent a heterogeneous collection of clinical strains, their genetic composition reflects the adaptations necessary for causing disease in humans. This large-scale genomic approach allows for the identification of genes and proteins critical for establishing infection in humans.

Proteins with low A/N (alanine-to-asparagine) indices are likely under strong selective pressure, suggesting their importance in maintaining cellular viability and supporting infection. The predicted essential proteins identified in this study could serve as potential drug targets aimed at disrupting *M. abscessus* survival within the host. Furthermore, understanding the functional roles of these proteins could provide insight into the unique metabolic and physiological adaptations that enable *M. abscessus* to thrive in the lung environment, where it predominantly causes chronic lung disease.

There are limitations to this study. We were unable to fully assemble the genomes of all isolates. Instead, we aligned protein sequences from the reference genome directly to the contigs. This approach excluded some isolates from our gene analysis, especially those with fragmented or incomplete assemblies. Additionally, we applied stringent threshold criteria to reduce the impact of large gene rearrangements, insertions, and deletions. Consequently, isolates from different *M. abscessus* subspecies with significant structural variation in certain genes may have been excluded, potentially leading to an underestimation of the true genetic diversity. As with any annotated genome, not all annotated proteins in ATCC 19977 have been experimentally validated. To generate reliable nucleotide and protein mutation maps, we included only high-quality gene sequences with unambiguous read data. Consequently, some sequences from certain isolates were excluded. However, the genome sequence collection available through NCBI was sufficiently large to generate high-quality sequence data for all target loci, ensuring robust statistical assessment and high confidence in the essentiality predictions. A larger genome collection would have permitted essentiality predictions on a larger number of genes/proteins especially those of the smaller sizes which we excluded from our analysis.

Future work should focus on experimentally validating these predicted essential proteins. Gene knockout studies and functional assays in relevant animal models of *M. abscessus* infection, which are becoming increasingly available, will be crucial for confirming the roles of these protein (41–44). Understanding the mechanistic basis of their function could reveal new vulnerabilities in *M. abscessus*, guiding the development of more effective treatments for *M. abscessus* lung disease.

## MATERIALS AND METHODS

### Packages and Softwares

Python (version 3.11.5), Biopython (version 1.84) (45), mappy (version 2.28) (46), shutup (version 0.2.0), scipy (version 1.13.1) (47), statsmodels (version 0.14.2) (48), pandas (version 2.2.2) (49), matplotlib (version 3.9.2) (50), openpyxl (version 3.1.5). R (version 4.4.2), reshape (version 1.4.4) (51), readxl (version 1.4.3) (52), ggplot2 (version 3.5.1) (53), seqinr (version 4.2.36) (54).

### Proteins with Atypical Amino Acid composition

Protein sequences from the *M. abscessus* ATCC 19977 (RefSeq ID GCF_000069185.1) were retrieved from a FASTA file using the SeqIO module of Biopython. The total proteome length and the counts for each amino acid were counted. The Global Proteome Amino Acid Distribution (GPAAD) in *M. abscessus* ATCC 19977 was then calculated by dividing the total count of each amino acid by the total proteome length.

For each protein, we calculated the number of amino acids and percentages of each amino acid. We used a binomial test for each amino acid in every protein to see if the numbers were different from the usual frequency of that amino acid in all proteins. Although a Bonferroni correction for multiple comparisons was considered, the raw p-values were retained for reporting due to the high baseline frequencies in the proteome (**Table S1**).

To mitigate the effect of statistical noise in shorter proteins, it was necessary to consider only proteins with a minimum of 100 amino acids when identifying outliers. For each amino acid, the proteins with the highest and lowest relative percentages were determined, and these extremes were subsequently summarized along with the overall proteome frequency in a separate Excel report.

Based on the outputs of the GPAAD and individual protein amino acid composition statistics, we identified a subset of protein outliers exhibiting significant deviations in the relative abundance of at least one amino acid compared to the overall proteome distribution (**Table 1**). In addition, protein length and associated p-values were taken into consideration to ensure that the selected proteins represent robust and biologically meaningful variations.

### Protein Regions with Atypical Amino Acid Composition

Protein sequences from the Mycobacterium abscessus ATCC 19977 were retrieved from a FASTA file using the SeqIO module of Biopython. First, the GPAAD was calculated, which served as the expected baseline distribution. Subsequently, each protein was segmented into overlapping regions of 100 amino acids with a 10-amino-acid step. For proteins shorter than 100 amino acids, the entire sequence was analyzed as a single region. In each region, the counted amino acid counts were compared to the expectation using the chi-square test.

For each protein, the three regions with the lowest p-values were identified and reported. The results, including the region’s location, p-value, and both observed and expected percentages for each amino acid, were then written to an Excel file for further evaluation.

A stringent significance threshold (p-value < 1×10⁻³⁰) was employed to identify a small subset of the most abnormal regions. For each protein, only the region with the lowest p-value was retained. The final results, which include the protein locus, protein length, enriched amino acid, and the corresponding abnormal region, are presented in Table 2.

### Proteins with Least Amino Acid Variety

First, the GPAAD was calculated, which served as the expected baseline distribution. For each individual protein, the presence or absence of each amino acid was assessed, and the amino acid variety was defined as 20 minus the number of missing amino acids, reflecting the number of amino acid usage in the protein. Then, a probability metric was calculated to quantify the likelihood of the observed absence of certain amino acids. This probability was computed under the assumption that the absence of an amino acid in a protein of given length follows a binomial model, where the chance of an amino acid being absent is derived from its proteome-wide frequency.

The resulting data, including protein ID, length, amino acid variety, list of missing amino acids, and the computed probability were saved into an excel file. A stringent significance threshold (p-value < 1×10⁻10) was employed to identify a small subset of the most abnormal proteins. All the selected proteins with the least amino acid variety were reported as Table 3.

### Alignment, Mutation Maps and Essential Gene Prediction

Reference gene sequences were first extracted from a FASTA file using a custom parser that retrieves the gene identifier, genomic coordinates, strand orientation, and nucleotide sequence. For genes located on the negative strand, we read the sequence in the complementary stain, so all genes have a uniform orientation, same as our WGS contigs. Mappy, a Python interface to minimap2, was used to align the data. For each sample, the reference was the genomic contigs, and the reference gene sequences were used as queries. Mappy generated a list of the reference contigs and used a seed-and-extend strategy to find potential alignment regions. This approach identifies short exact matches (seeds) between the query and reference and then connects them. Then, a process called dynamic programming was used to find and correctly place any mismatches, insertions, and deletions.

We set thresholds to avoid long range rearrangements, insertions and deletions, to check the mapping quality. These criteria were 95% coverage and 92% identity. For each high-quality alignment, the aligned sequence was extracted, and alignment metrics (coverage, identity, CIGAR string, alignment position, and mapping quality) were recorded. These metrics were written into an alignment report, and the final set of aligned sequences was saved in the corresponding files for each gene for further analysis.

We initialized MAFFT to align the reference genes and their matches among 430 isolates found in the previous step. The first sequence in each alignment file served as the reference, which was derived from the *M. abscessus* ATCC 19977 genome. When a gene was identified on the complementary strand, its sequence was reverse complemented to ensure correct alignment.

The aligned sequences were then processed with a custom mapping function that relates each position in the aligned (gapped) sequence to its corresponding position in the original, ungapped sequence. This mapping allowed us to accurately identify and count mutation events at specific nucleotide positions by comparing each isolate sequence to the reference.

The aligned gene sequences were translated into protein sequences using translation table 11 for bacteria. Then, the same procedure was applied to the protein alignments to determine mutation frequencies at the amino acid level.

Finally, mutation frequencies were calculated as the fraction of isolates showing a mutation at each position. The resulting mutation data includes both gene and protein mutation frequencies and the ratio of protein to gene mutations. Mutation maps were drawn to visualize the mutation data (**Figure 2**).

## AUTHOR CONTRIBUTIONS

GC: methodology, study design, investigation, data analysis and interpretation, manuscript preparation. RAH: data analysis and interpretation, manuscript preparation. BR: data analysis and interpretation, manuscript preparation. NN. Methodology and data interpretation. GL: study conception, study design, project administration, data interpretation, manuscript preparation, and funding acquisition.

