## Supplemental Information for "Predicting *Mycobacterium abscessus* proteins with atypical amino acid composition essential for human infections"

**FIGURES**

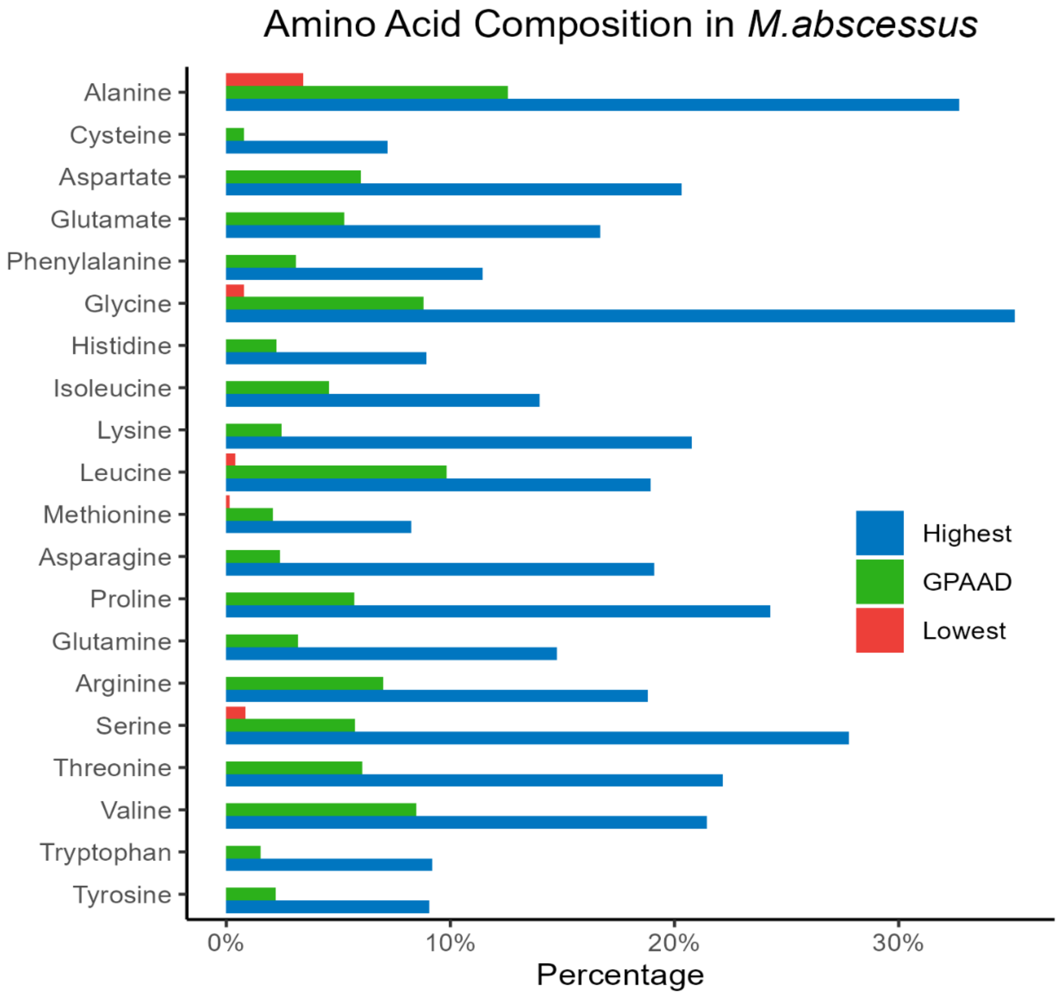


**Figure S1**. Distribution of each amino acid in *Mycobacterium abscessus* ATCC 19977 proteome, denoted by Global Proteome Amino Acid Distribution (GPAAD). Also shown are the lowest (RED) and highest (BLUE) percentage composition of an amino acid in M. abscessus proteins.

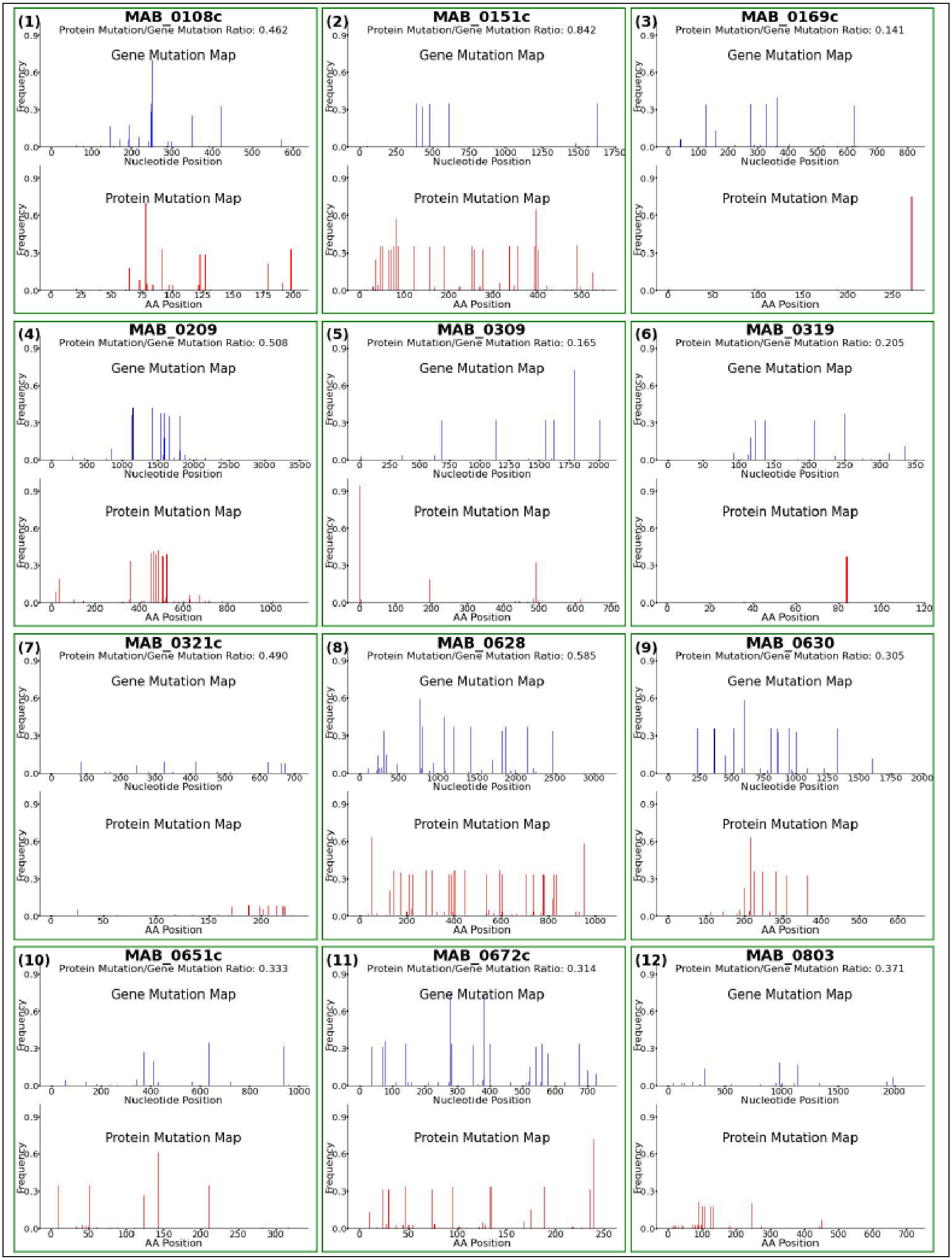

(1)
MAB_0108c
Protein Mutation/Gene Mutation Ratio: 0.462
(2)
MAB_0151c
Protein Mutation/Gene Mutation Ratio: 0.842
(3)
MAB_0169c
Protein Mutation/Gene Mutation Ratio: 0.141
(4)
MAB_0209
Protein Mutation/Gene Mutation Ratio: 0.508
(5)
MAB_0309
Protein Mutation/Gene Mutation Ratio: 0.165
(6)
MAB_0319
Protein Mutation/Gene Mutation Ratio: 0.205
(7)
MAB_0321c
Protein Mutation/Gene Mutation Ratio: 0.490
(8)
MAB_0628
Protein Mutation/Gene Mutation Ratio: 0.585
(9)
MAB_0630
Protein Mutation/Gene Mutation Ratio: 0.305
(10)
MAB_0651c
Protein Mutation/Gene Mutation Ratio: 0.333
(11)
MAB_0672c
Protein Mutation/Gene Mutation Ratio: 0.314
(12)
MAB_0803
Protein Mutation/Gene Mutation Ratio: 0.371
Gene Mutation Map
Protein Mutation Map
Frequency
Nucleotide Position
AA Position

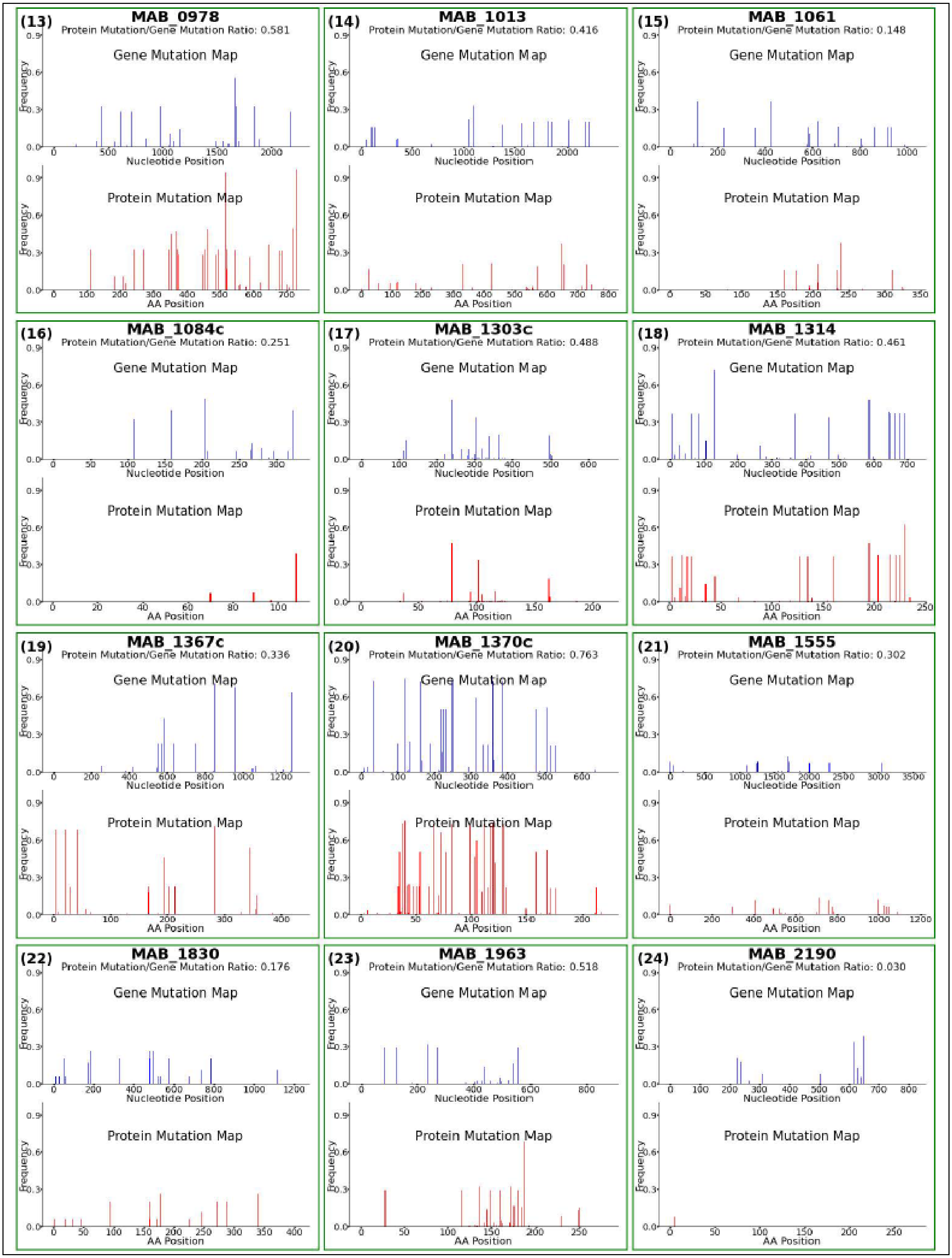
(13)
MAB_0978
Protein Mutation/Gene Mutation Ratio: 0.581
(14)
MAB_1013
Protein Mutation/Gene Mutation Ratio: 0.416
(15)
MAB_1061
Protein Mutation/Gene Mutation Ratio: 0.148
(16)
MAB_1084c
Protein Mutation/Gene Mutation Ratio: 0.251
(17)
MAB_1303c
Protein Mutation/Gene Mutation Ratio: 0.488
(18)
MAB_1314
Protein Mutation/Gene Mutation Ratio: 0.461
(19)
MAB_1367c
Protein Mutation/Gene Mutation Ratio: 0.336
(20)
MAB_1370c
Protein Mutation/Gene Mutation Ratio: 0.763
(21)
MAB_1555
Protein Mutation/Gene Mutation Ratio: 0.302
(22)
MAB_1830
Protein Mutation/Gene Mutation Ratio: 0.176
(23)
MAB_1963
Protein Mutation/Gene Mutation Ratio: 0.518
(24)
MAB_2190
Protein Mutation/Gene Mutation Ratio: 0.030
Gene Mutation Map
Protein Mutation Map
Frequency
Nucleotide Position
AA Position

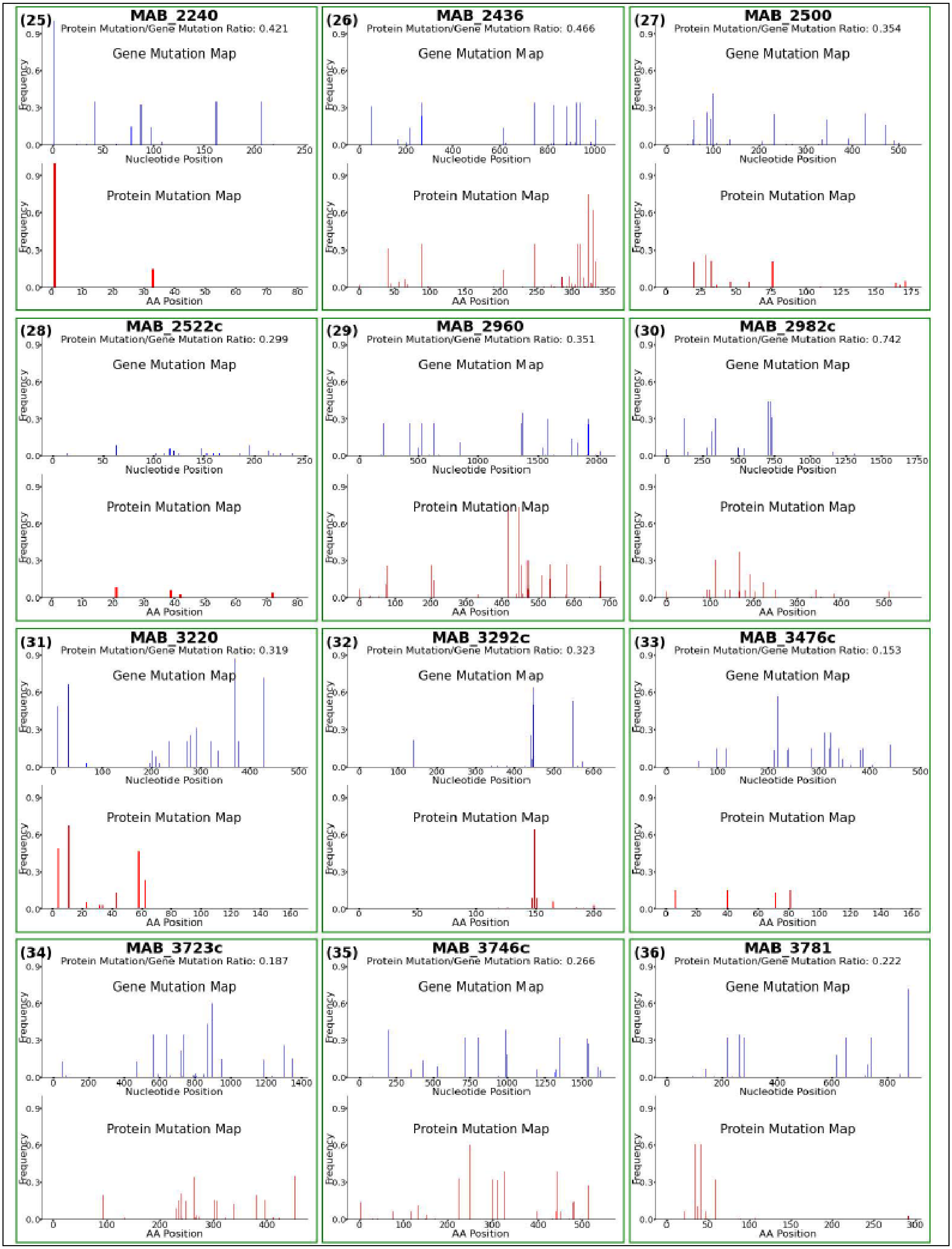
(25) MAB_2240
Protein Mutation/Gene Mutation Ratio: 0.421
(26) MAB_2436
Protein Mutation/Gene Mutation Ratio: 0.466
(27) MAB_2500
Protein Mutation/Gene Mutation Ratio: 0.354
(28) MAB_2522c
Protein Mutation/Gene Mutation Ratio: 0.299
(29) MAB_2960
Protein Mutation/Gene Mutation Ratio: 0.351
(30) MAB_2982c
Protein Mutation/Gene Mutation Ratio: 0.742
(31) MAB_3220
Protein Mutation/Gene Mutation Ratio: 0.319
(32) MAB_3292c
Protein Mutation/Gene Mutation Ratio: 0.323
(33) MAB_3476c
Protein Mutation/Gene Mutation Ratio: 0.153
(34) MAB_3723c
Protein Mutation/Gene Mutation Ratio: 0.187
(35) MAB_3746c
Protein Mutation/Gene Mutation Ratio: 0.266
(36) MAB_3781
Protein Mutation/Gene Mutation Ratio: 0.222
Gene Mutation Map
Protein Mutation Map
Frequency
Nucleotide Position
AA Position

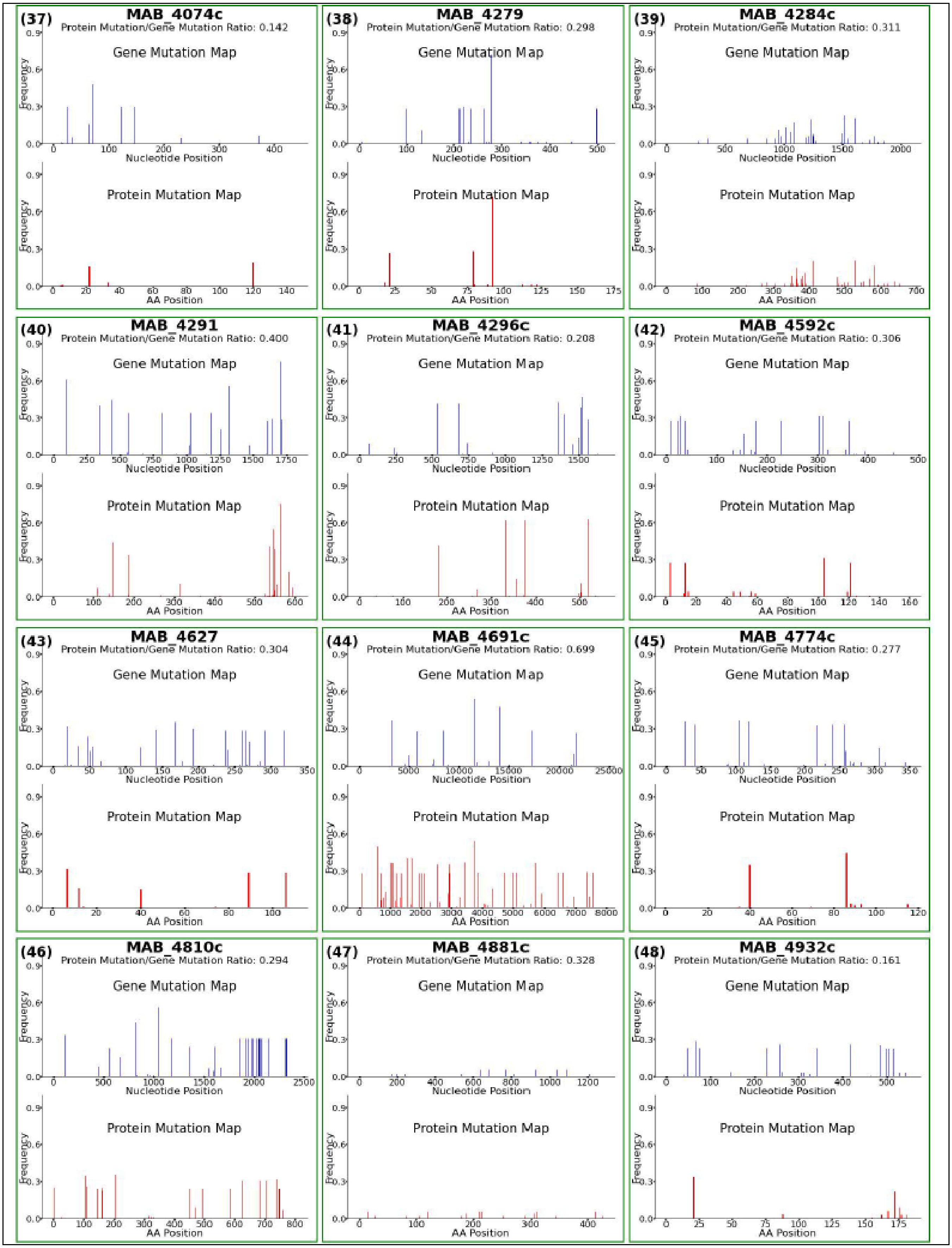
(37) MAB_4074c
Protein Mutation/Gene Mutation Ratio: 0.142
(38) MAB_4279
Protein Mutation/Gene Mutation Ratio: 0.298
(39) MAB_4284c
Protein Mutation/Gene Mutation Ratio: 0.311
(40) MAB_4291
Protein Mutation/Gene Mutation Ratio: 0.400
(41) MAB_4296c
Protein Mutation/Gene Mutation Ratio: 0.208
(42) MAB_4592c
Protein Mutation/Gene Mutation Ratio: 0.306
(43) MAB_4627
Protein Mutation/Gene Mutation Ratio: 0.304
(44) MAB_4691c
Protein Mutation/Gene Mutation Ratio: 0.699
(45) MAB_4774c
Protein Mutation/Gene Mutation Ratio: 0.277
(46) MAB_4810c
Protein Mutation/Gene Mutation Ratio: 0.294
(47) MAB_4881c
Protein Mutation/Gene Mutation Ratio: 0.328
(48) MAB_4932c
Protein Mutation/Gene Mutation Ratio: 0.161
Gene Mutation Map
Protein Mutation Map
Frequency
Nucleotide Position
AA Position

**Figure S2**. **Mutation maps that illustrate the evolution of *Mab* genes/proteins that are unlikely to be essential for causing disease in humans**. Each panel shows the mutation profile of a specific gene and corresponding protein, highlighting the nucleotide and amino acid positions and the frequency of mutations in patient-derived isolates.

### TABLES

**Table S1**. This is a large worksheet that is included as a separate file.

| Gene/Protein ID | Gene/Protein Name | Nucleotide Mutation Count | Amino Acid Mutation Count | Amino Acid Mutation/Nucleotide Mutation Ratio |
|---|---|---|---|---|
| MAB_0001 | DnaA | 2372 | 128 | 0.054 |
| MAB_0002 | DnaN | 4439 | 77 | 0.017 |
| MAB_3770c | RpoA | 2680 | 2 | 0.001 |
| MAB_3868c | RpoC | 6737 | 222 | 0.033 |
| MAB_3869c | RpoB | 7461 | 807 | 0.108 |
| | | | | **Mean = 0.043** |

**Table S2**. List of five known essential proteins in *M. abscessus*. The number of nucleotide and amino acid mutations in the corresponding genes/proteins in patient derived isolates are shown. The last column lists the ratio of amino acid mutation to nucleotide mutation (A/N).

| Protein ID | Gene Mutation Count | Protein Mutation Count | Amino Acid Mutation/Nucleotide Mutation (A/N) |
|---|---|---|---|
| MAB_0030 | 8 | 8 | 1.000 |
| MAB_0108c | 2248 | 1039 | 0.462 |
| MAB_0151c | 6259 | 5267 | 0.842 |
| MAB_0169c | 2183 | 308 | 0.141 |
| MAB_0209 | 6405 | 3251 | 0.508 |
| MAB_0309 | 7291 | 1202 | 0.165 |
| MAB_0319 | 742 | 152 | 0.205 |
| MAB_0321c | 382 | 187 | 0.490 |
| MAB_0628 | 20271 | 11852 | 0.585 |
| MAB_0630 | 9357 | 2850 | 0.305 |
| MAB_0651c | 2442 | 812 | 0.333 |
| MAB_0672c | 5087 | 1598 | 0.314 |
| MAB_0775 | 45 | 25 | 0.556 |
| MAB_0803 | 2523 | 937 | 0.371 |
| MAB_0978 | 8832 | 5131 | 0.581 |
| MAB_1013 | 9400 | 3912 | 0.416 |
| MAB_1061 | 4132 | 612 | 0.148 |
| MAB_1084c | 869 | 218 | 0.251 |
| MAB_1303c | 1079 | 527 | 0.488 |
| MAB_1314 | 4568 | 2104 | 0.461 |
| MAB_1367c | 7267 | 2440 | 0.336 |
| MAB_1370c | 7544 | 5756 | 0.763 |
| MAB_1555 | 1521 | 459 | 0.302 |
| MAB_1830 | 1864 | 328 | 0.176 |
| MAB_1963 | 2794 | 1447 | 0.518 |
| MAB_2190 | 99 | 3 | 0.030 |
| MAB_2240 | 1097 | 462 | 0.421 |
| MAB_2436 | 3808 | 1775 | 0.466 |
| MAB_2500 | 1175 | 416 | 0.354 |
| MAB_2522c | 177 | 53 | 0.299 |
| MAB_2960 | 6956 | 2439 | 0.351 |
| MAB_2982c | 1878 | 1393 | 0.742 |
| MAB_3220 | 254 | 81 | 0.319 |
| MAB_3292c | 1131 | 365 | 0.323 |
| MAB_3476c | 255 | 39 | 0.153 |
| MAB_3723c | 6741 | 1261 | 0.187 |
| MAB_3746c | 7519 | 2000 | 0.266 |
| MAB_3781 | 3287 | 729 | 0.222 |
| MAB_4074c | 1099 | 156 | 0.142 |
| MAB_4279 | 342 | 102 | 0.298 |
| MAB_4284c | 1732 | 539 | 0.311 |
| MAB_4291 | 9263 | 3702 | 0.400 |
| MAB_4296c | 6298 | 1311 | 0.208 |
| MAB_4592c | 1800 | 551 | 0.306 |
| MAB_4627 | 1444 | 439 | 0.304 |
| MAB_4691c | 95943 | 67046 | 0.699 |
| MAB_4774c | 1309 | 362 | 0.277 |
| MAB_4810c | 9035 | 2653 | 0.294 |
| MAB_4881c | 244 | 80 | 0.328 |
| MAB_4932c | 2097 | 337 | 0.161 |
| MAB_5000c | 2 | 2 | 1.000 |

**Table S3**. List of proteins in *M. abscessus* with atypical amino acid distribution and their amino acid mutation / nucleotide mutation ratios. The number of nucleotide and amino acid mutations in the corresponding genes/proteins in patient derived isolates are shown. The last column lists the ratio of amino acid mutation to nucleotide mutation (A/N).
